# Effects of different environmental intervention durations on the intestinal mucosal barrier and the brain-gut axis in rats with colorectal cancer

**DOI:** 10.1101/2022.03.17.484783

**Authors:** Liu Dun, Chen Mei-Jing, Huang Si-Ting, Yu Xin-Yuan, Wu Yu-Xuan

## Abstract

**Objectives:** Enriched environment (EE) is a promising strategy to protect the intestinal mucosal barrier and regulate brain-gut peptide expression. However, the optimal enriched environment intervention duration is unknown. Here, different EE intervention durations were applied to assess the optimal intervention duration in rats with colorectal cancer.

**Methods:** We used a rat model of 1,2-dimethylhydrazine-induced colorectal cancer. Rats were housed in an EE for 0, 2, 4 or 8 weeks. The intestinal mucosa and serum TNF-α, IL-6, IL-10, DAO, ATP, CRF, and occludin levels and bacterial translocation (BT) were measured, and the intestinal mucosa morphology was evaluated.

**Results:** Eight-week EE intervention was more beneficial to the intestinal mucosal mechanical barrier than 2-week or 4-week intervention (P<0.05). There was a significant different between the 4-week and 8-week groups on BT (P=0.049). However, which intervention duration had the greatest advantages in intestinal mucosa and serum inflammatory factor regulation was not determined. There were no significant differences in the effects of different EE intervention durations on BT or brain intestinal peptide levels among the other groups (P>0.05).

**Conclusions:** The effect of an 8-week environmental intervention duration on the intestinal mucosal barrier was better than that of 2-week and 4-week durations overall, but the effect of different environmental intervention durations on brain-gut peptide levels was not obvious. In the future, we can further explore the molecular biological mechanism of the effect of different EE intervention durations on the intestinal mucosal barrier and analyze the effect of an EE on other brain-gut peptides.

## INTRODUCTION

Enriched environment(EE) is a paradigm in which animals are introduced to novel, complex, and stimulating surroundings that promote structural changes in the brain and enhance learning and memory performance in rodents.^1^ In an enriched environment, mice or rats are placed in larger cages containing multiple physical and social stimuli, which allow the animals to explore, exercise, and interact with each other.^2^ Recent studies have investigated the role of EEs in cancer treatment and prognosis.^2–5^ Several studies have indicated that EEs can regulate the intestinal microenvironment and enhance intestinal immunity,^6–11^ thereby protecting the intestinal mucosal barrier. Previous studies have shown that the combination of the three factors of enriched environment, social interaction and cognitive stimulation played a role in improving the levels of intestinal ghrelin, serum IL-6, intestinal mucosal TNF-α and IL-6, and hypothalamic corticotrophin-releasing factor (CRF). The effect of combining social interaction and cognitive stimulation on ghrelin levels in the hypothalamus, TNF-α levels in serum and IL-10 levels in serum and the intestinal mucosa was better than the effect of the combination of the three factors. However, the combination of the three factors was better than the combination of social interaction and cognitive stimulation in improving muscle layer thickness and occludin levels. Among the univariate factors, cognitive stimuli improved TNF-α levels in serum and IL-10 levels in the intestinal mucosa more than the other two factors.^12^ In addition, EEs can regulate the expression of brain-gut peptides,^13–16^ thereby regulating gastrointestinal function. Therefore, an EE is beneficial for the protection of the intestinal mucosa.

However, in the above studies, the EE intervention duration ranged from 2 to 18 weeks. The optimal duration of the EE intervention is not well known. Therefore, in this study, we used different durations of EE intervention in colorectal cancer (CRC) rats to analyze the optimal EE intervention duration.

## MATERIAL AND METHODS

### Animals

All procedures were approved by the Laboratory of Animal Welfare and Ethics Committee of our university (certificate number 2016-06) and were performed in accordance with the National Institutes of Health guidelines for the treatment of animals. We obtained 60 four-week-old male Sprague–Dawley rats weighing approximately 150–160 g each from the Animal Experimental Center of Fujian Medical University. The rats were housed in ventilated cages containing four to five rats and soft shavings in an air-conditioned room (22±1°C, 50–60% humidity, 12-h light-dark cycle), and the rats were fed ad libitum (20–25% protein, 5–10% fat, and 3–5% crude fiber diet). The food was prepared and mixed according to the guidelines set by the Association of Analytical Communities. Prior to the experiments, the rats were fed adaptively based on the criteria described above for two weeks. Rats that did not exhibit loss of agility or loss of teeth were deemed fit for the investigation.

### Generation of 1,2-dimethylhydrazine-induced tumors

From six weeks of age, the rats were subcutaneously injected with 1,2-dimethylhydrazine (DMH; Sigma Chemical Co., USA) (concentration 2%, 20 mg/kg, pH 6.5) once per week for 21 weeks. DMH is highly carcinogenic and has high organ specificity. DMH has been used in many studies to model CRC in rats.^17^

### Animal housing procedures

Twenty-one weeks after DMH injection, all rats were examined by ultrasonography (Esaote, MylabClassC, Italy; probe frequency, 18–22 MHz). Tumor formation occurred in all 48 rats, and these animals were then divided into four groups of 12 rats each based on stratified randomization according to their weights. The first group was the blank group. The EE intervention durations in the second, third and fourth groups were 2 weeks, 4 weeks and 8 weeks, respectively. Previous studies have shown an elevated level of prostaglandin E2 in the colonic mucosa of patients with CRC.^18^ Prostaglandin E2 is associated with malignant tumor progression and intestinal barrier function.^19^ A higher BMI is associated with a higher level of prostaglandin E2.^20^ Therefore, in this study, weight was used as a baseline measure of tumor progression and bowel function in cancer.

For the EE conditions, large cages (109 × 79 × 41 cm) containing twelve rats were used. An EE was established as described in the relevant literature.^21–24^ Stimulatory objects were placed in the cages. The number of stimulatory objects was approximately 1–2 per rat, and the objects included huts made of wood, walking wheels made of plastic with a diameter of 21 centimeters, transparent labyrinth tunnels made of acrylic with a diameter of 13 centimeters and various wooden toys. All of the objects were harmless to the rats. Items that had been destroyed by the rats were replaced periodically. In addition, the positions of the stimuli, water, and food in the cages were changed twice weekly.

### Western blotting

Prior to tissue sample collection, all rats were anesthetized and then sacrificed. A piece of colon tissue (approximately 100 mg) located 2 cm from the end of the cecum was collected and washed with saline. Tissues were then immediately placed in an Eppendorf tube and stored at −80°C. Approximately 100 mg of tissue was treated with 200 μl of Protein Seeker Mammalian Cell Lysis Solution (GenDEPOT, USA). The tissue was ground in ice-cold water, shaken at 1200 rpm for 5 s and then centrifuged at 12,000–16,000 rpm for 5 min. The supernatants were collected and stored at −80°C prior to use. Extracted proteins were analyzed by sodium dodecyl sulfate– polyacrylamide gel electrophoresis and then transferred onto a polyvinylidene fluoride membrane (Pall). Western blotting was performed using a polyclonal anti-ghrelin primary antibody (1:250; Abcam, UK). The membranes were then incubated with horseradish peroxidase-conjugated secondary antibodies (1:8000, ZB-2301, Zhongshan Company, Beijing) for 1 h. The immunoreactive proteins were detected using the ECL-Plus Western Blotting Detection System (Amersham Life Sciences, Braunschweig, Germany).

### Measurement of TNF-α, IL-6, IL-10, and CRF levels

Rat intestines (approximately 400 mg of tissue per sample) were clipped and washed with saline. Then, the samples were cut into slices, homogenized using a Dounce homogenizer (WHEATON, USA), and centrifuged at 4°C and 10,000 rpm for 30 min. The supernatants from each fraction were collected and stored at −80°C. Brain tissues were isolated in an ice bath, and the hypothalami were separated and rapidly placed in Eppendorf tubes. The tissues were then frozen in liquid nitrogen for 5 min and stored at −80°C. For serum samples, blood was drawn from the inner canthus vein. Whole blood was incubated at 4°C for 24 h and then centrifuged at 10,000 rpm for 10 min. Serum was isolated from whole blood using a liquid transfer gun and stored at −80°C. All samples were then thawed, and the levels of TNF-α, IL-6, IL-10, and CRF were determined by enzyme-linked immunosorbent assays (ELISAs) according to the manufacturer’s instructions.

### Detection of bacterial translocation

Mesenteric lymph nodes, livers, and spleens were collected under sterile conditions. To each specimen, 1 ml of cold saline was added. Then, the samples were ground using a mortar and pestle, and 0.5 ml of each sample was incubated with medium containing eosin methylene blue agar at 37°C for 48 h. The rosin acid contained in the medium inhibited the growth of gram-positive bacteria but had no inhibitory effect on the growth of gram-negative bacteria. Escherichia coli is a gram-negative bacterium that is capable of lactose decomposition and forms blue colonies on this medium, with the size of a single colony being approximately 2 mm.^25^ The number of bacterial colonies was counted, and the number of CFUs per gram of tissue (CFUs/g) was calculated. In this study, culture plates harboring more than five colonies indicated a positive result.^26^

### Intestinal mucosal morphology

Rat intestines were isolated and fixed with 10% formaldehyde, dehydrated, and embedded in paraffin. Then, 5-μm-thick sections were cut, dewaxed with xylene, hydrated with an alcohol gradient, and stained with hematoxylin for 1 min. The samples were then washed with phosphate-buffered saline (PBS) and stained with eosin for 15 s before being rapidly dehydrated with an alcohol gradient. Finally, the sections were treated with xylene, mounted in neutral gum, and viewed with a light microscope. Hematoxylin and eosin (HE) staining results were evaluated using Image-Pro Plus 6.0, and the groups were compared.

### Immunohistochemical detection of occludin

Rat intestines were fixed in 10% formalin for 24 h and embedded in paraffin. Then, 5-μm thick sections were cut, mounted on slides, and incubated with an anti-occludin (1:120, Thermo, USA) for 2 h at 37°C. The slides were then washed three times with PBS and incubated with a goat anti-rabbit secondary antibody (Maixin Biological Technology Development Co., Ltd., Fuzhou, China) for 30 min. The slides were washed three times with PBS and developed using diaminobenzidine color development solution (Fuzhou Maixin Biotechnology Development Co., Ltd.) for 5 min. The slides were then stained with hematoxylin for 1 min, washed with PBS, dehydrated with an alcohol gradient, treated with xylene, mounted with neutral gum, and viewed with a light microscope (Nikon SMZ645, Japan). Immunohistochemical staining results were evaluated using Image-Pro Plus 6.0, and between-group comparisons were conducted. The investigator who analyzed all immunohistochemically stained slides was blinded to the group allocation of each sample. The expression levels of occludin were analyzed as described in the relevant literature.[36]

### Statistical analysis

Bacterial translocation (BT) measured in various organs is presented as a percentage. The other indices are presented as the means ± standard deviations. Differences between groups were assessed using two-factor analysis of variance (ANOVA) with a post hoc Bonferroni pairwise comparison. Weight differences were assessed using two-way classification repeated ANOVA. For no normally distributed data, a Kruskal–Wallis test with post hoc Mann–Whitney U test for pairwise comparison was performed. P<0.05 was considered significant. An adjusted significance level of P<0.01 was used for post hoc pairwise comparisons. All statistical analyses were performed using SPSS 24.0 statistics (SPSS Inc., Chicago, IL, USA).

## RESULTS

### Microscopic morphology of the intestinal mucosa

#### Villus length

Samples were collected from the rat intestines, and pathology methods were used for analysis. There was a significant effect among the different groups on small intestinal villus length (F=74.920, P=0.000). The small intestinal villus length was not significantly different between the 4-week group and the 8-week group (P=0.473). There were significant differences among the other groups (Figure 1).

**Figure 1.**
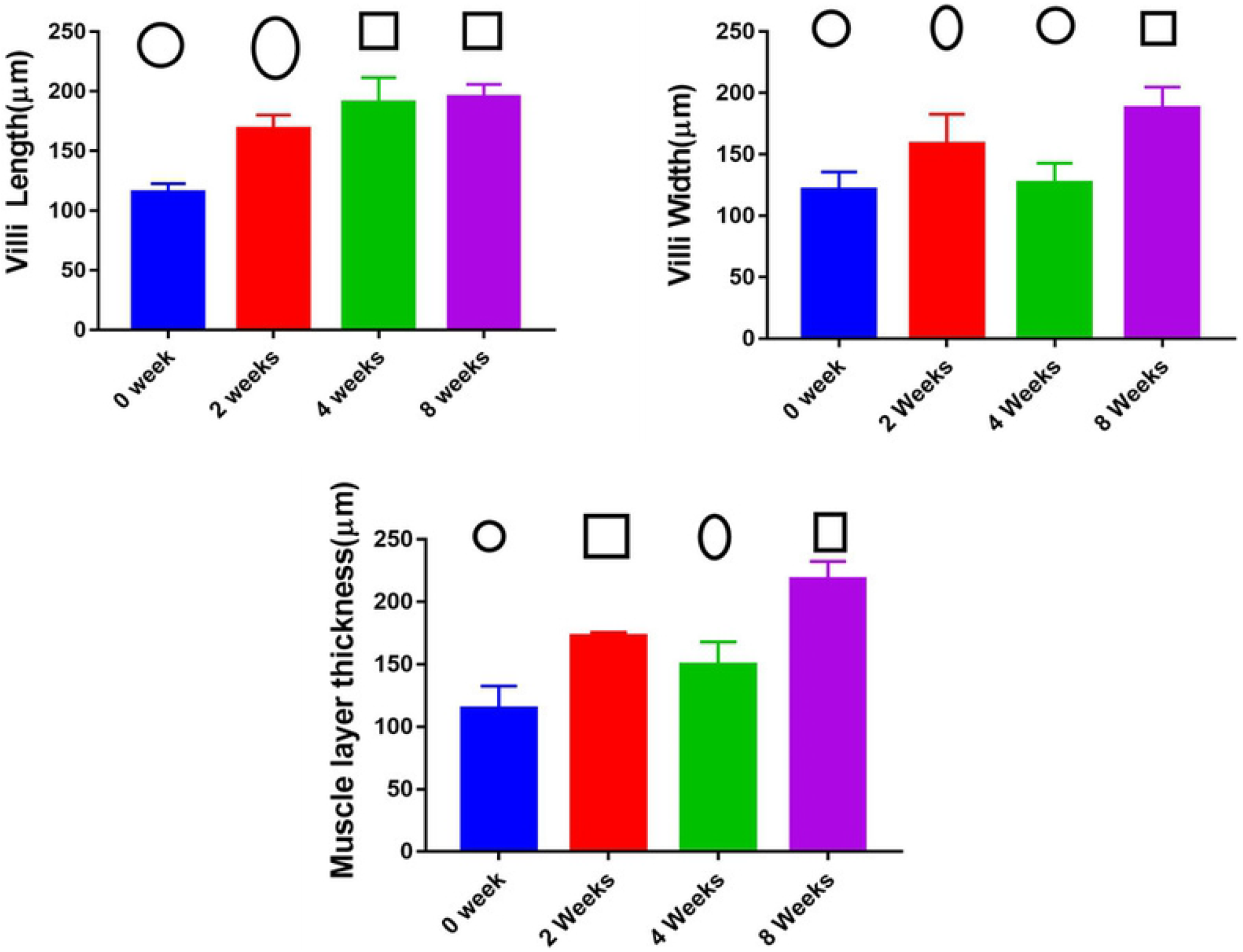

#### Villus width

There was a significant effect among the different groups on small intestinal villus width (F=25.979, P=0.000). The small intestinal villus width was not significantly different between the blank group and the 4-week group (P=0.463). There were significant differences among the other groups (Figure 1).

#### Muscle layer thickness

There was a significant effect among the different groups on small intestinal muscle layer thickness (F=20.516, P=0.000). There was no significant difference between the 2-week and 4-week groups, but there were significant differences among the other groups (Figure 1).

### Immunohistochemical detection of occludin

There was a significant effect among the different groups on intestinal occludin (F=20.516, P=0.000). The intestinal occludin staining was not significantly different between the 2-week group and the 4-week group (P=0.353). There were significant differences among the other groups (Table 1).

### Serum and intestinal mucosa levels of TNF-α, IL-6, and IL-10

Serum samples were collected, and ELISAs were used for detection. There was a significant effect among the different groups on the serum levels of IL-10 (F=13.951, P=0.000). The serum levels of IL-10 were not significantly different between the 2-week group and the 4-week group (P=0.228) or between the blank group and the 8-week group (P=0.123). There were differences among the other groups (Table 2, Figure 2).

**Figure 2.**
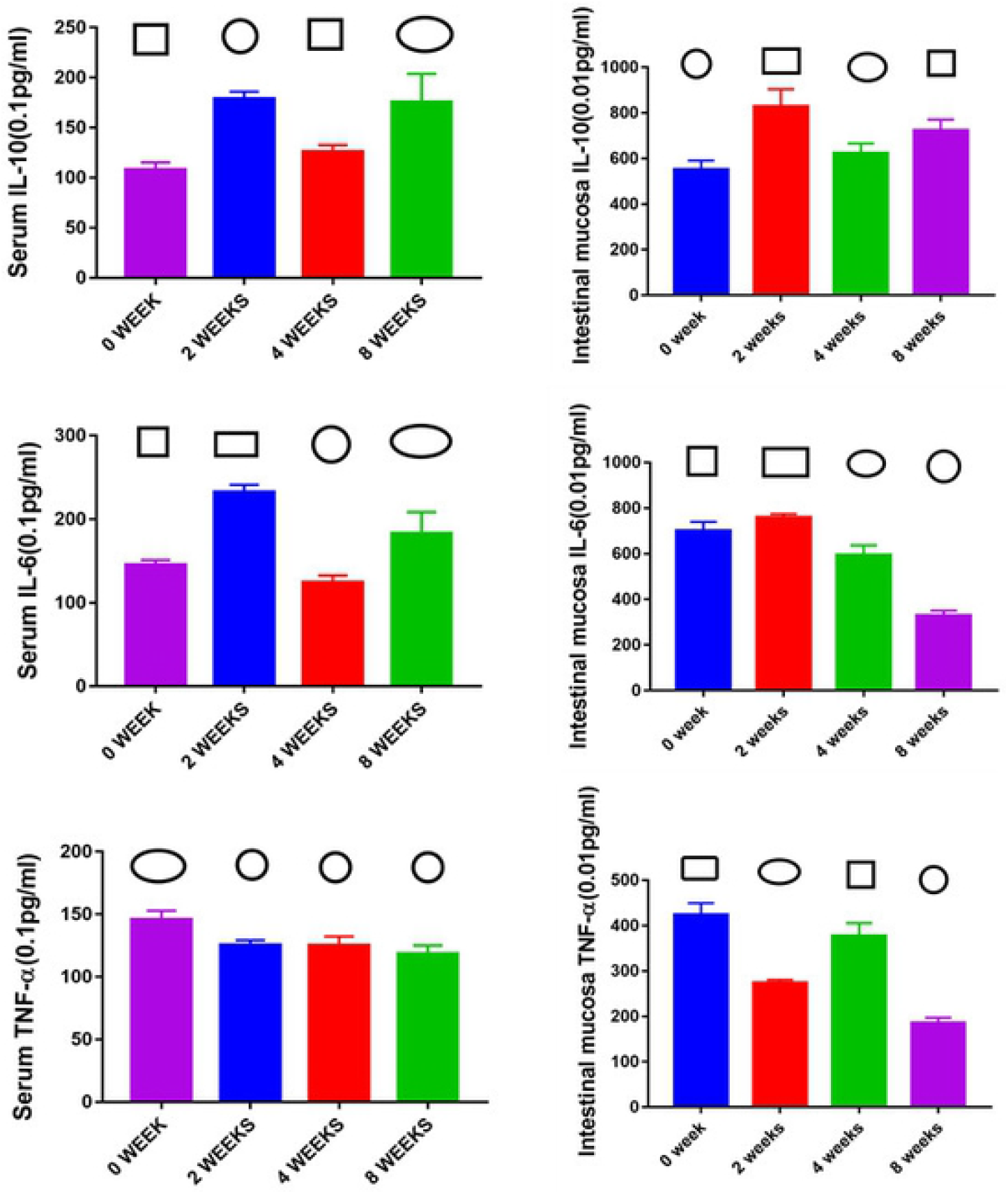

There was a significant effect among the different groups on the serum levels of IL-6 (F=52.119, P=0.000). The serum levels of IL-10 were not significantly different between the blank group and the 4-week group (P=0.261) or between the 2-week group and the 4-week group (P=0.545). There were differences among the other groups (Table 2, Figure 2).

There was a significant effect among the different groups on the serum levels of TNF-α (F=60.345, P=0.000). The serum levels of TNF-α were not significantly different between the 2-week group and 4-week group (P=0.853), the 4-week group and 8-week group (P=0.608) and the 2-week group and 8-week group (P=0.647). There were differences among the other groups (Table 2, Figure 2).

There was a significant effect among the different groups on the intestinal mucosa levels of IL-6 (F=306.547, P=0.000). The intestinal mucosa levels of IL-6 were statistically significantly different among the different groups (Table 2, Figure 2). There was a significant effect among the different groups on the intestinal mucosa levels of IL-10 (F=58.191, P=0.000). The intestinal mucosa levels of IL-10 were significantly different among the different groups (Table 2, Figure 2).

There was a significant effect among the different groups on the intestinal mucosa levels of TNF-α (F=251.150, P=0.000). The intestinal mucosa levels of TNF-α were significantly different among the different groups (Table 2, Figure 2).

### BT ratios in the three groups

The results indicated that BT occurred in 20 out of 36 tissues in the two-week group, 23 out of 36 tissues in the 4-week group and 8 out of 24 tissues in the 8-week group. There was a significant effect between the 4-week group and the 8-week group on BT (P=0.049). There were no significant differences among the other groups.

### Brain-gut peptide levels in rats with CRC

#### Serum CRF levels in rats with CRC

Serum samples were collected, and an ELISA was used for detection. There was no significant effect among the different groups on the serum levels of CRF (F=0.282, P=0.756).

## DISCUSSION

Some studies have shown that an EE not only influences brain structure and function ^27^ but also significantly inhibits tumor growth in colon cancer.^28^ A previous study on EE intervention for 8 weeks in rats with CRC also showed that an EE could increase brain–gut peptide expression (especially ghrelin secretion) and enhance intestinal mucosal immunologic functions, thus ameliorating intestinal dysfunction and maintaining the integrity of the intestinal mucosal barrier. However, the best and most effective intervention durations are still not known. Therefore, this study used different intervention durations for CRC rats to analyze the optimal EE intervention duration for rats with CRC to lay a foundation for the study of nondrug interventions for clinical cancer patients.

### Intestinal mucosal mechanical barrier

Intestinal epithelial cells and tight junctions (TJs) between intestinal epithelial cells form the structural basis of the intestinal mucosal mechanical barrier. TJ barrier function can be affected by changes in the distribution of specific TJ proteins and/or their expression levels. The intestinal epithelial transmembrane binding protein occludin is a transmembrane protein that is one of the primary TJ proteins among the zonula occludens proteins.^29–31^ Therefore, occludin levels and the length, thickness, and muscular thickness of intestinal epithelial villi can affect the intestinal mucosal mechanical barrier to a certain extent.

The results of this study show that in general, the effect of 8-week EE intervention weeks is better than that of 2- and 4-week interventions. The intervention effect at 2 and 4 weeks was better than that observed in the blank group. However, it is uncertain whether the 2-week intervention time or the 4-week intervention time is better. Therefore, an EE is helpful to maintain the intestinal mechanical barrier, and the effect of longer intervention times is better than that of shorter intervention times. The effect with less than 4 weeks of intervention may not be superior.

### Intestinal mucosal immune barrier

Cytokines are the major regulators of mucosal immunity and are important for the intestinal immune defense response.^32^ Cancer patients generally exhibit changes in cytokine levels, which greatly affect metabolism and immunity in the body. There are two primary types of cytokines: (1) those that promote the inflammatory reaction, such as TNF-α, IL-1, and IL-6, and (2) those involved in the suppression of inflammatory response factors, such as IL-4 and IL-10.^33^

Our results showed that an EE can adjust IL-10, IL-6, and TNF-α levels to exert beneficial effects on the body. However, there were still differences among groups with different intervention times. Serum IL-6 levels in the 4-week group were lower than those in the other groups. The levels of serum IL-10 in the 2-week and 8-week groups were higher than those in the 4-week group, and the levels of serum IL-10 in the 3 intervention groups were higher than those in the blank group. The serum TNF-α level in the 2-week and 4-week groups was higher than that in the 8-week group, and the serum TNF-α level in the 3 intervention groups was lower than that in the blank group. Therefore, overall, an EE helps regulate inflammatory factors, thereby protecting the intestinal immune barrier. The intervention effect at different times was better than that in the blank group, and the intervention effect at 8 weeks was slightly better than that at 2 and 4 weeks. However, the effect of different intervention times on the intestinal mucosal immune barrier needs to be further studied for additional inflammatory factors to yield a clearer conclusion.

### Intestinal mucosal biological barrier

BT refers to the translocation of intestinal bacteria from the intestinal lumen to the mesentery or other organs. Normally, intestinal BT does not occur easily owing to tight intestinal junctions. However, BT in the intestinal tract increases during bacterial pathogenesis or during periods of stress when the mucosal epithelium is damaged. Therefore, BT can be used to evaluate the permeability of the intestinal mucosal barrier.^34^ In this study, although there was a difference between only the 4-week and 8-week groups, overall, the regulatory effect of the 8-week intervention on BT was better than that of the 2-week and 4-week interventions. Therefore, after sufficient EE intervention time, the intervention can regulate the intestinal mucosal biological barrier. These results suggest that adequate cognitive, psychological, sport and other nondrug intervention measures are conducive to the improvement in intestinal mucosal biological barrier function. However, the specific mechanism needs further study.

### Brain-gut peptide

CRF is the primary mediator that allows the central nervous system to participate in stress responses. Under stress conditions such as disease, CRF can be overexpressed to regulate gastrointestinal motility, secretion and sensation through the hypothalamic-pituitary-adrenal (HPA) axis.^35^ Regarding the regulation of CRF expression, different EE intervention durations had no significant effect on brain intestinal peptide levels. This result may be related to the intensity of the intervention or other factors. Thus, the effect of EE intervention on the intestinal mucosa may not necessarily occur through the regulation of the brain-intestine axis. Therefore, the effects of different intervention durations on the brain-intestine axis need to be further investigated.

## Conclusions

The effect of an 8-week environmental intervention duration on the intestinal mucosal barrier was better than that of a 2-week or 4-week duration overall, but the effect of different environmental intervention durations on brain-gut peptide levels was not obvious. In the future, we can further explore the molecular biological mechanism of the effect of different EE intervention durations on the intestinal mucosal barrier and analyze the effect of an EE on other brain-gut peptides. This study lays a foundation for clinical nondrug intervention research for cancer.

## Acknowledgements

We thank Mr. Xiang-xin Wu for technical assistant. We also grateful for Prof. Cai-hua Huang for her advice on EE housing set up.

## Ethical Approval and Consent to participate

Male specific pathogen-free (SPF) Sprague-Dawley (SD) rats were purchased from the Animal Experiment Center of Fujian Medical University. The rats were purchased under the license number SCXK (Fujian) 2016-0002. After purchase, the rats were reared at the Animal Experiment Center of Fujian Medical University under the license number SYXK (Fujian) 2016-0006. All procedures involving animals were performed in accordance with the ethical standards of the institution or practice at which the studies were conducted (Fujian Medical University, 2016-06).

## Availability of supporting data

Others can be able to replicate and build upon the authors’ published claims. Authors can make materials, data, code, and associated protocols promptly available to readers without undue qualifications. Any restrictions on the availability of materials or information must be disclosed to the editors at the time of submission. Any restrictions must also be disclosed in the submitted manuscript.

## Conflict of Interest

All authors participated in the design and interpretation of the studies, analysis of the data and review of the manuscript. Chen Mei-jing conducted the experiments, Liu Dun supplied crucial reagents and animals, and Liu Dun, Huang Si-Ting, Yu Xin-Yuan, and Wu Yu-Xuan wrote the manuscript. The authors declare that they have no conflicts of interest.

## Financial support

This study was funded in full by the Fujian Natural Science Foundation Project (grant no. 2021J01789) and the Fujian Provincial Joint Innovation Project of Science and Technology Department (grant no. 2018Y9101).

## Notes

### Competing Interest Statement

The authors have declared no competing interest.

